# Development of a microbiome based health score for non-invasive monitoring of farmed Atlantic salmon (*Salmo salar*)

**DOI:** 10.64898/2026.02.27.708536

**Authors:** Luis E. León, Catalina Lorca, Francisco Fuentes, Agustín Piña, María Ignacia Ortuzar, Diego Gutierrez, Juan A. Ugalde, Alejandro Bisquertt

**Affiliations:** Codebreaker Biosciences, Puerto Varas, Chile; Center for Bioinformatics and Integrative Biology, Facutad de Ciencias de la Vida, Universidad Andres Bello, Santiago, Chile; SENTINET: Surveillance, Epidemiology, and New Technologies for Infectious Emerging Threats, Santiago, Chile

**Keywords:** Health score, microbiome, *Salmo salar*, aquaculture, machine learning, Atlantic salmon, precision aquaculture

## Abstract

The sustainability of Atlantic salmon (*Salmo salar*) farming is threatened by infectious diseases, environmental stressors, feed limitations, and regulatory or economic constraints. Although current health monitoring is improving with AI-powered camera systems that analyze behavior and nutrition, these tools typically identify stress responses rather than early signs of disease states. Because the microbial community undergoes successional reassembly in response to physiological disruptions before host barriers are breached, the microbiome offers a proactive early warning approach. In this study, a comparative cohort design was employed using a total of 171 individuals (n= 85 “healthy”; n=86 “lesioned”) collected from a commercial marine facility and classified based on external clinical signs. The microbiome of multiple body sites (gills, skin, urogenital pore, and mucosa) from healthy and lesioned salmon were profiled using 16S rRNA amplicon sequencing. No differences in alpha diversity were observed between tissues and conditions. However, beta diversity was significantly different in clinical status, and the interaction of tissue with the status. Conversely, the mucosa and urogenital microbiomes were compositionally similar to each other, as were the gill and skin microbiomes, suggesting that urogenital swabs could serve as a non-invasive proxy for gut microbiome profiling, and skin for gill microbiomes. Several supervised models were trained on these profiles and used to classify salmon status with high accuracy. Based on these data, two Salmon Microbiome Health Score were developed that accurately differentiated the two cohorts. These scores are proposed as a novel biomarker, enabling proactive health management in aquaculture and complementing emerging technological monitoring systems.

## Introduction

Aquaculture represents one of the fastest growing production sectors globally, playing an increasingly critical role in meeting the nutritional needs of a growing human population (Froehlich et al., 2018; Troell et al., 2014). Atlantic salmon (*Salmo salar*) farming is a central component of this industry, with global production reaching significant figures each year and holding substantial economic importance. However, the sustainability and profitability of salmon aquaculture remain fundamentally constrained by multifactorial health challenges, including but not limited to infectious diseases, environmental stressors, and management-related conditions.

The result of these multifactorial health challenges is a complex landscape of contemporary production issues. For instance, complex gill disease (CGD) has emerged as one of the most significant health challenges in salmon farming (Mitchell and Rodger, 2011). Bacterial infections such as tenacibaculosis, caused by species of Tenacibaculum genus (Mabrok et al., 2023; Spilsberg et al., 2022; Wynne et al., 2020), and salmonid rickettsial septicaemia (SRS), caused by *Piscirickettsia salmonis*, continue to impose major economic and welfare impacts. SRS, in particular, remains one of the most devastating bacterial diseases in Chilean aquaculture, characterized by systemic infection, high mortality, and limited vaccine efficacy (Evensen, 2016; Rozas-Serri, 2022; Rozas-Serri et al., 2024).

Traditional farm health management relies on visual identification of clinical manifestations such as cutaneous lesions, behavioral changes, or gill pathology (Noble et al., 2026, 2012). This reactive approach is suboptimal, as disease is typically advanced by the time signs manifest. In response, the industry has transitioned toward “precision aquaculture”, emphasizing early detection through real-time monitoring of biological and environmental parameters (Burke et al., 2025). Artificial intelligence technologies, particularly computer vision systems monitoring behavior, feeding patterns, and biomass, support this transformation. However, these tools primarily detect existing stress or pathologies, creating a need for approaches capable of assessing biological health at earlier stages.

The host-associated microbiome is a highly sensitive indicator of physiological state and health (Clinton et al., 2024). These microbial communities perform essential functions including nutrient acquisition, immune maturation, epithelial barrier maintenance, and pathogen exclusion (Auclert et al., 2024; Wang et al., 2018; Xiong et al., 2019). Research in several organisms including salmon indicates that certain microbial signatures correlate with host health status irrespective of disease etiology, potentially allowing identification of biomarkers prior to the appearance of clinical signs (Bozzi et al., 2021; Hajjo et al., 2022; Wang et al., 2023).

Foundational studies have mapped the spatial organization of the salmon gut microbiome (Gajardo et al., 2016), characterized the effects of environmental transitions on intestinal communities (Dehler et al., 2017), and demonstrated that interpopulation variation reflects both environmental and genetic diversity . The salmon microbiome is now recognized as a hologenomic component with significant implications for aquaculture management (Limborg et al., 2018).

In teleost fish, the intestinal and branchial microbiomes are of primary importance. The gut microbiome is central to digestive physiology, nutrient metabolism, and systemic immunity. The gill microbiome constitutes the primary interface with waterborne pathogens and environmental stressors, with tissue and mucus microbiomes changing during disease episodes (Wynne et al., 2020). Monitoring the microbiota of these tissues provides direct insight into early health trajectories. Importantly, microbial community responses to stress often follow the Anna Karenina principle, whereby dysbiotic individuals exhibit greater compositional variability than their healthy counterparts (Zaneveld et al., 2017), a pattern that has been documented in fish during disease outbreaks.

Despite this potential, routine microbiome monitoring faces a critical constraint: sampling informative niches requires invasive procedures incompatible with production-scale aquaculture. Gill sampling requires individual capture and handling, inducing acute stress and logistic challenges (Clinton et al., 2021). Gut characterization relies overwhelmingly on lethal sampling. These requirements render invasive sampling unsuitable for longitudinal monitoring of valuable stock.

To address this limitation, we tested the hypothesis that easily accessible, non-invasive microbiomes can serve as proxies for the health status reflected by internal sites. Skin and branchial epithelium are distinct mucosal surfaces in continuous contact with the same aquatic environment. We hypothesized these sites harbor compositionally similar communities exhibiting coordinated responses to pathogens, water quality, and stressors. Similarly, the urogenital opening’s anatomical proximity to the anus suggests the intestinal microbial community continuously seeds the urogenital tract, resulting in compositional similarity. Evidence from other fish species supports these proxy relationships; for example, external mucus microbiomes have been shown to reflect internal conditions such as chronic enteritis in yellowtail kingfish (Legrand et al., 2017).

The aims of this study were threefold: (1) to validate the compositional concordance between non-invasive (skin, urogenital pore) and invasive (gill, mucosa) microbiome sampling sites; (2) to develop and benchmark machine learning models using non-invasive sample data to classify salmon health status; and (3) to formulate these models into two quantitative, non-invasive health scores for practical implementation in proactive aquaculture health management.

## Materials and methods

A total of 171 Atlantic salmon (Salmo salar), were collected from a commercial marine aquaculture facility. All fish were from the same production cohort to minimize confounding effects. With the assistance of on-site veterinarians, the fish were classified into one of two groups: “healthy” (n =85) or “lesioned” (n = 86). Fish were assigned to the lesioned cohort if they presented one or more distinct clinical signs associated with common salmonid diseases. These signs included skin lesions or ulcers, exophthalmos, or visible pale gills. Fish in the healthy cohort showed no external signs of pathology. This binary classification provided the basis for all subsequent comparative analysis.

### Sample collection

Following classification, the fish were euthanized via immersion in an overdose of MS-222 (tricaine methanesulfonate) buffered with sodium bicarbonate. Sample collection was conducted during routine sanitary evaluations under established Standard Operating Procedures (SOPs) and under strict veterinary supervision. Since the sampling involved non-experimental, routine diagnostic procedures and no additional interventions beyond standard farm health assessments were performed, formal ethical committee approval was not required. All procedures were carried out in compliance with animal welfare protocols to ensure the ethical handling of the fish . To obtain a holistic profile of the salmon-associated microbiome, samples were aseptically collected from four anatomical niches. A sterile swab was passed firmly three times over the gill arches to sample the gills, and a separate swab was rubbed within a 4 cm^2^ area on the lateral surface posterior to the dorsal fin for the skin sample. A third swab was gently inserted approximately 0.5 cm into the urogenital opening and rotated to collect the corresponding sample. Finally, for the digestive tract, a segment of the posterior intestine was carefully excised, and the intestinal mucus (or mucosa) was collected by gently scraping the inner surface.Immediately after collection, samples were placed into a tube containing a preservation solution, placed on ice, and stored at -80°C until DNA extraction.

### DNA extraction and library preparation

Total genomic DNA was extracted from each sample using the ZymoBIOMICS DNA MagBead Kit following the manufacturer’s specifications. Each DNA extract was quantified using a Qubit fluorometer for accurate measurement of double-stranded DNA concentration. DNA quality and amplifiability were assessed based on qPCR amplification curves, including evaluation of amplification efficiency and overall curve performance.

High-throughput sequencing of the 16s ribosomal RNA gene was conducted at Codebreaker Biosciences facilities in Puerto Varas, Chile. Amplification was performed using primers 341f (CCTACGGGNGGCWGCAG) and 806r (GACTACHVGGGTATCTAATCC) (Klindworth et al., 2013), targeting the V3-V4 region of the 16S rRNA gene, using the Quick-16S™ NGS Library Preparation Kit (Zymo Research, CA). The negative control utilized was DNase/RNase-Free Water (Zymo Research, CA). Following amplification, PCR products were quantified using qPCR fluorescence readings and pooled based on equal molarity. The pooled library underwent purification with the Select-a-Size DNA Clean & Concentrator™ (Zymo Research, CA) and was subsequently quantified using Qsep1 (Bioptic) and Qubit (Thermo). Sequencing was performed on an Illumina® NextSeq 1000™ platform with a P1 reagent kit (600 cycles), configured for 2 × 300 bp paired-end reads and incorporating a 10% PhiX spike-in control. To account for potential reagent or environmental contamination critical for low-biomass sites such as skin and gills negative controls (extraction and PCR blanks) were sequenced alongside samples.

### Bioinformatic analysis

The raw FASTQ files were processed using DADA2 (version 1.22) (Callahan et al., 2016) which infers exact Amplicon Sequence Variants (ASVs). To account for variable quality profiles across the sequencing run, a dynamic truncation strategy was employed, identifying the coordinate where the median quality score dropped below a threshold of 25. Following initial quality filtering and trimming, error rates were learned and reads were further filtered using a maximum expected error (*maxEE*) threshold of 2 for both forward and reverse reads. Paired-end reads were merged with a minimum overlap of 20 bp, and chimeric sequences were identified and removed using the *removeBimeraDenovo* consensus method. The DADA2 workflow has been chosen for its ability to infer exact Amplicon Sequence Variants (ASVs), which provides single-nucleotide resolution. The pipeline included quality filtering and trimming, error rate learning, sample inference, merging, and chimera removal, resulting in a final ASV abundance table.

Contamination control was performed through rigorous inspection of the negative extraction and PCR controls sequenced alongside the samples. ASVs present in negative controls with a relative abundance exceeding 1% or those identified as common laboratory contaminants (e.g., *Pseudomonadota, Bacillota*) were manually flagged and removed from the dataset. Samples with fewer than 20,000 reads were excluded to ensure sufficient sequencing depth. Furthermore, a 10% prevalence threshold within each tissue type and a minimum requirement of 10 reads in at least two samples served as an additional stringency filter to eliminate low-frequency noise and potential environmental artifacts typical of low-biomass swabs.

Taxonomic identity was assigned to each ASV using the DADA2 implementation of the Naive Bayes classifier using 16s rRNA gene reference sequences from the Genome Taxonomy Database (GTDB) (Parks et al., 2022) version R226. For assignments reaching the species level, the classification reflects the best genomic match. Due to the inherent resolution limits of the 16s rRNA marker, these species-level assignments were treated as presumptive. Alpha diversity was estimated using the Shannon diversity index (*H*) and Simpson’s dominance index (*D*). Prior to centered log ratio (CLR) transformation, zeros were imputed using Bayesian multiplicative replacement via the *zCompositions* R package (*cmultRepl* function with geometric bayesian multiplicative (GBM) method, frac=0.65). This approach ensures that the compositional integrity of the microbial signatures is maintained. Subsequently, compositional beta diversity was evaluated by applying a CLR transformation to the imputed abundance table, followed by Principal Component Analysis (PCA) on the transformed data using euclidean distance. All analyses were conducted using the phyloseq R package (McMurdie and Holmes, 2013).

### Multivariate statistical analysis of microbial communities

Permutational Multivariate Analysis of Variance (PERMANOVA) was employed to evaluate differences in microbial community composition using the adonis2 function in the vegan R package. The global model assessed the effects of anatomical site (Source), clinical status (Status), and their interaction using Aitchison distances (Euclidean distances on CLR transformed data). To account for the paired sampling design and mitigate inter-individual variability, permutations (n=999) were strictly constrained within each individual fish.

Subsequently, multivariate homogeneity of group dispersions was evaluated using the betadisper function to validate PERMANOVA assumptions and characterize community stability across health statuses (Anderson, 2006). To rigorously validate the use of non invasive proxies, the concordance between invasive and non-invasive pairs (Mucosa vs. Urogenital pore and Gill vs. Skin) was assessed through a dual validation approach: (i) Procrustes analysis (protest function) to compare Principal Component Analysis (PCA) ordination configurations, and (ii) Mantel tests using Spearman’s rank correlation to quantify the strength of distance matrix coupling. Significance for all tests was determined through 999 permutations (Peres-Neto and Jackson, 2001).

### Classification models

To identify the most informative taxonomic resolution, a combinatorial search was performed across multiple three-level aggregation schemes (e.g., Class-Order-Species). This phase of the modeling was exploratory, focusing on comparing the predictive signals of different feature sets rather than producing final validated classifiers. L1-regularized logistic regression and random forests were applied within a nested, group aware crossvalidation framework to rank these combinations by their relative predictive capability (Topçuoğlu et al., 2020). Subsequently, the most predictive taxonomic sets identified through this process were used in the next steps.

A comprehensive classifier benchmark was implemented to predict health status from microbiome profiles, systematically comparing Random Forest, XGBoost, LightGBM, Support Vector Machines (SVM), Logistic Regression, K-Nearest Neighbors (KNN), and Gaussian Naive Bayes. To ensure unbiased performance estimation across sea cages, a nested cross-validation framework with group blocking was utilized (Roberts et al., 2017). This structure comprised an outer 2-fold stratified group cross-validation for final performance evaluation and an inner 5-fold cross-validation for hyperparameter tuning. Group blocking by sea cage prevented data leakage from samples within the same sea cage appearing in both training and test sets. This design guaranteed that the test set data remained completely independent of the optimization process, preventing optimistic bias in the performance metrics. A model-specific preprocessing strategy was adopted: for tree-based ensemble models (Random Forest, XGBoost, LightGBM), ASV counts were normalized using Total Sum Scaling (TSS), while for linear and distance-based models (SVM, KNN, Logistic Regression, Naive Bayes), a CLR transformation was applied separately to each taxonomy level to respect the compositional nature of mixed-level taxonomic data, followed by z-score standardization and PCA, retaining 75% of the variance for dimensionality reduction. All transformations were exclusively fitted on the training data of each respective fold. Hyperparameter optimization was performed via grid search within the inner loop. Model performance was assessed using accuracy, F1-score, and AUC-ROC, reported as mean ± standard deviation across the outer folds. To identify the most robust diagnostic model, a multi-criteria decision analysis (MCDA) was employed. Rather than relying on a single metric, models were ranked based on a weighted composite score comprising: AUC (30%), overall Accuracy (15%), Balanced Accuracy (15%), Matthews Correlation Coefficient (15%), Consistency/Stability (5%), Brier Score for calibration (15%), and Cohen’s Kappa (5%). All metrics were appropriately scaled, with the Brier Score inverted so that lower values contributed to a higher final score. This systematic approach ensures that the selected best model optimizes not only for predictive power but also for reliability, class-balance robustness, and calibration quality. Youden’s J statistic for optimal threshold determination was also calculated.

### Salmon microbiome health scores

Inspired by the development of quantitative health indices in human microbiome research, such as the Gut Microbiome Wellness Index (GMWI2) (Chang et al., 2024), the objective was to translate the complex outputs of the top-performing models into intuitive, site-specific metrics. Two distinct indices were formulated based on the predictive outputs of the logistic regression models: a Gut Health Score (GHS), derived from urogenital pore samples, and a Skin Health Score (SHS), derived from gill samples.

Both scores were defined as the model-predicted probability that an individual fish belongs to the lesioned cohort, yielding a continuous value ranging from 0 to 1 (healthy to lesioned). To validate the biological relevance of these metrics, the distribution of scores was compared between healthy and lesioned cohorts using Mann-Whitney U tests.

## Results

After performing quality control and filtering, a final dataset of 599 samples was retained from an initial pool of 676. Sample distribution remained balanced across most anatomical sites and health statuses, though a reduction in healthy skin samples was observed probably due to low microbial biomass (Table 1). The loss of healthy skin samples likely reflects the lower microbial load on intact skin surfaces, resulting in more samples failing to meet the 20,000 read sequencing depth threshold. This differential attrition should be considered when interpreting the Skin Health Score. To mitigate potential classification bias arising from this imbalance, the machine learning framework utilized stratified group cross-validation and weighted scoring metrics, ensuring that the model’s performance reflects biological signals rather than sample frequency disparities.

**Table 1.** Summary of dataset dimensions highlighting sample retention. Numbers represent individual samples (n) and the corresponding percentage categorized by anatomical site and clinical status (Healthy vs. Lesioned). The filtered stage represents the final dataset used for all downstream multivariate analyses, including PCA, Mantel tests, and machine learning classifiers.

| Tissue | Status | Raw | Filtered |
| --- | --- | --- | --- |
| Gill | Healthy | 85 | 79 |
|  | Lesioned | 84 | 84 |
| Mucosa | Healthy | 81 | 79 |
|  | Lesioned | 84 | 79 |
| Skin | Healthy | 85 | 35 |
|  | Lesioned | 86 | 74 |
| Urogenital pore | Healthy | 84 | 83 |
|  | Lesioned | 87 | 86 |
| **Total** | Healthy | 335 | 276 |
|  | Lesioned | 341 | 323 |

### Microbial composition and diversity

The analysis of 16S rRNA gene sequences revealed distinct microbial community compositions driven by the interaction between anatomical niche and health status. Taxonomic profiles highlight the compositional differences across the four sampled body sites (Figure 1).

**Figure 1.**
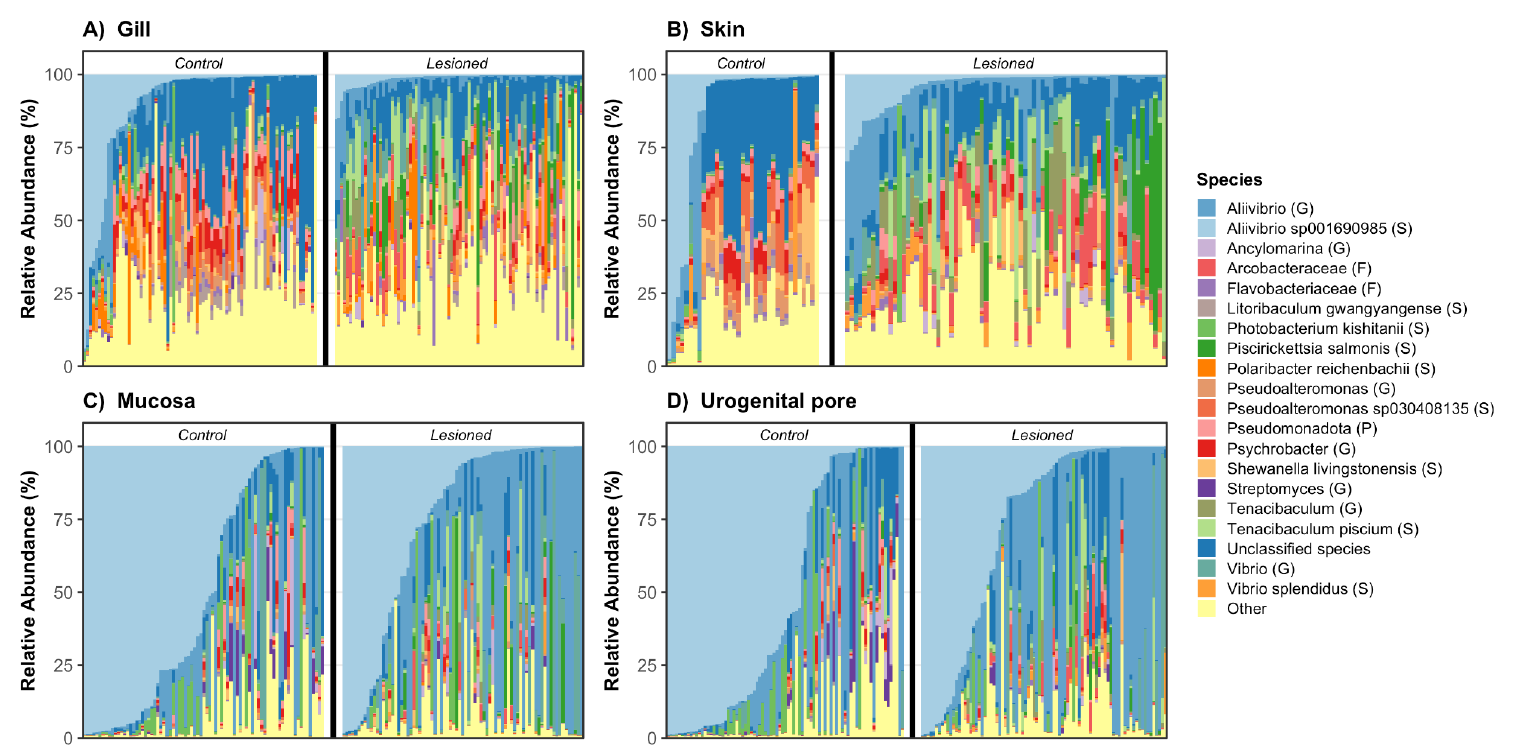
Relative abundance (%) of the top 20 taxa at the lowest taxonomic level across individual samples. Each stacked bar represents a single sample; panels are faceted by Source (Gill, Skin, Mucosa, and Urogenital pore), and Status (Healthy and Lesioned). Colors indicate taxa as shown in the legend. Taxa names include the lowest taxonomic rank reached, using abbreviations in parentheses.

It is important to note that taxonomic assignments reaching the species level (e.g., *Piscirickettsia salmonis* or *Vibrio splendidus*) are based on the best genomic match within the GTDB database for the V3-V4 16S rRNA region. Given the inherent resolution limits of this marker, these assignments should be considered presumptive. The biological interpretation of the health score primarily relies on broader successional patterns rather than the definitive presence of specific pathogens.

In contrast to the clear taxonomic shifts, analysis of microbial alpha diversity (Shannon and Simpson indices) revealed no significant differences between healthy and lesioned individuals within any of the four sampling zones (Table 2). This indicates that while the specific members of the community changed during disease, the overall complexity, richness, and evenness remained stable.

**Table 2.** Summary of microbial alpha diversity metrics by anatomical site and health status. Values are presented as mean ± standard deviation (SD). Comparison between Healthy and Lesioned individuals was performed using the Wilcoxon rank-sum test. P-values were adjusted for multiple comparisons within each metric using the False Discovery Rate (FDR) method.

| Metrics | Source | Healthy | Lesioned | p.adj |
| --- | --- | --- | --- | --- |
| Shannon | Gill | 3.13 ± 0.83 | 3.23 ± 0.62 | 0.78 |
| Shannon | Skin | 2.86 ± 0.87 | 3.06 ± 0.67 | 0.47 |
| Shannon | Mucosa | 1.63 ± 0.91 | 1.71 ± 0.88 | 0.47 |
| Shannon | Urogenital pore | 1.60 ± 0.98 | 1.78 ± 0.87 | 0.13 |
| Simpson | Gill | 0.17 ± 0.14 | 0.12 ± 0.10 | 0.35 |
| Simpson | Skin | 0.22 ± 0.16 | 0.14 ± 0.12 | 0.07 |
| Simpson | Mucosa | 0.44 ± 0.22 | 0.39 ± 0.21 | 0.26 |
| Simpson | Urogenital pore | 0.45 ± 0.22 | 0.38 ± 0.20 | 0.10 |

### Beta diversity and community concordance

To assess the factors driving community structure, a PERMANOVA was performed. The PERMANOVA analysis revealed that clinical status significantly reshapes the salmon microbiome (R^2^ = 0.038, F = 26.44, p = 0.006), with a significant interaction effect indicating that the microbial response to lesions is niche-specific (R^2^ = 0.018, F = 4.20, p = 0.001). Post-hoc dispersion analysis (betadisper) confirmed that these differences were driven by compositional centroid shifts in the urogenital pore (p = 0.466), Mucosa (p = 0.823), and Skin (p = 0.213), whereas the gills exhibited significantly higher dispersion in lesioned fish (F = 4.83, p = 0.029), suggesting a breakdown of microbial homeostasis.

Spatial coupling between anatomical interfaces was statistically robust. Procrustes analysis demonstrated strong concordance for both Skin-Gill (*r*_*Proc*_ = 0.842, *p* = 0.001) and Mucosa-Urogenital pore (*r*_*Pro*c_ = 0.701, *p* = 0.001) pairs. These findings were independently corroborated by Mantel tests, which showed significant distance matrix correlations for Skin-Gill (*r*_*M*_ = 0.52, *p* = 0.001, n = 106 pairs) and Mucosa-Urogenital (*r*_*M*_ = 0.54, p=0.001, n=154 pairs). These results support the use of external swabs as biological assessing internal health status.

**Figure 2.**
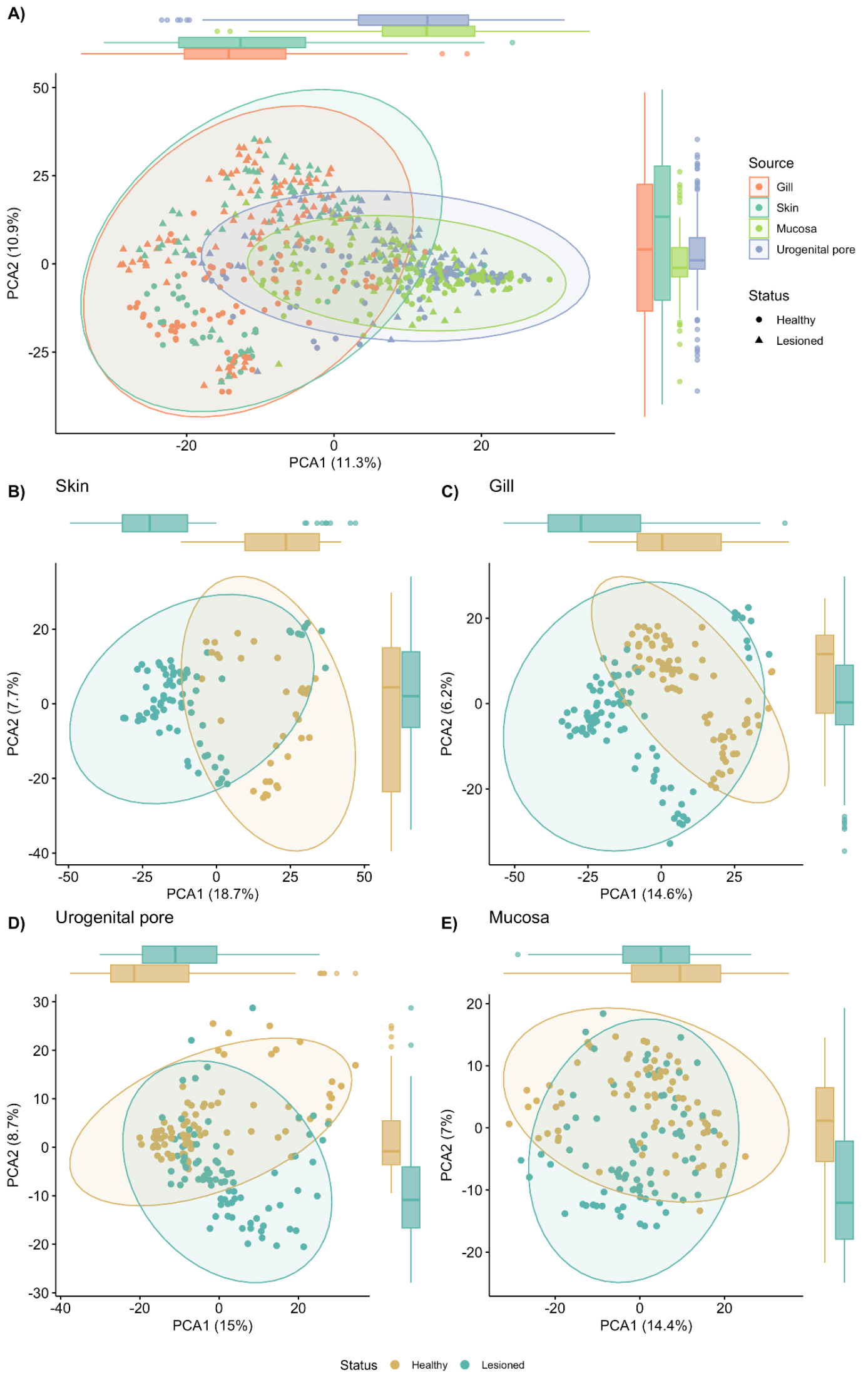
Principal Components Analysis (PCA) plot. PCA plot of euclidean distances of CLR transformed abundances, illustrating the beta diversity of microbial communities. Each point represents a single sample, A) colored by body site and shaped by health status, B) Skin, C) Gill, D) Urogenital pore, and E) Mucosa colored by health status.

### Machine learning classification

To identify the most effective algorithm for predicting salmon health status, seven different machine learning classifiers were trained and evaluated for skin and urogenital pore. The performance of all models is shown in Table 3. The results indicate that Logistic Regression achieved the highest overall predictive performance for both skin and for urogenital pore. Accordingly, these models were selected for all subsequent biomarker identification and health score formulation.

**Table 3.** Systematic selection of top-performing diagnostic models using Multi-Criteria Decision Analysis (MCDA). Models are ranked by a Weighted Score as described in the methods section. The Brier Score evaluates the accuracy of predicted probabilities (lower values indicate better calibration). Bold entries signify the primary models selected for health index development.

| Source | Rank | Model | Weighted Score | AUC mean | Accuracy mean | Balanced Accuracy mean | MCC mean | Brier Score |
| --- | --- | --- | --- | --- | --- | --- | --- | --- |
| Skin | 1 | Logistic Regression | 0.78 | 0.97 | 0.87 | 0.80 | 0.70 | 0.09 |
|  | 2 | Random Forest | 0.65 | 0.89 | 0.79 | 0.68 | 0.44 | 0.15 |
|  | 3 | XGBoost | 0.63 | 0.81 | 0.75 | 0.70 | 0.44 | 0.18 |
| Urogenital pore | 1 | Logistic Regression | 0.72 | 0.87 | 0.78 | 0.78 | 0.58 | 0.18 |
|  | 2 | SVM | 0.64 | 0.82 | 0.75 | 0.76 | 0.53 | 0.23 |
|  | 3 | KNN | 0.63 | 0.80 | 0.76 | 0.76 | 0.53 | 0.20 |

**Figure 3.**
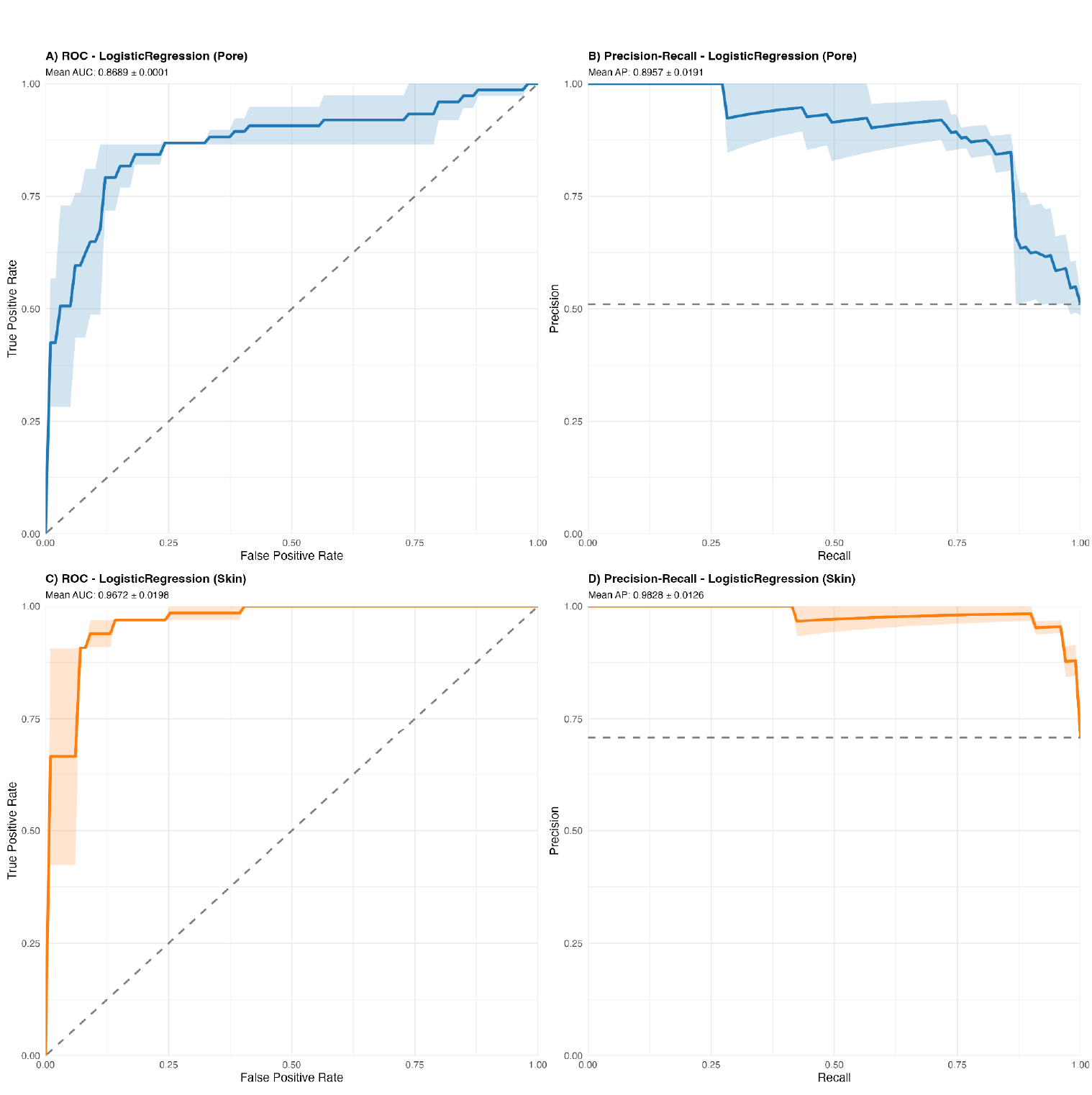
Discriminative power of non-invasive microbial proxies. Receiver Operating Characteristic (ROC) and Precision-Recall (PR) curves for the top-performing diagnostic models. Panels (A) and (B) show the ROC and PR curves, respectively, for the Urogenital pore using a Logistic Regression classifier. Panels (C) and (D) show the ROC and PR curves for the Skin using a Logistic Regression classifier. Shaded areas represent the 95% confidence intervals across 5-fold cross-validation. The high Area Under the Curve (AUC) values in both the ROC and PR spaces demonstrate that these non-invasive sites provide a robust and precise signal for health status classification, with minimal trade-off between sensitivity and specificity.

**Figure 4.**
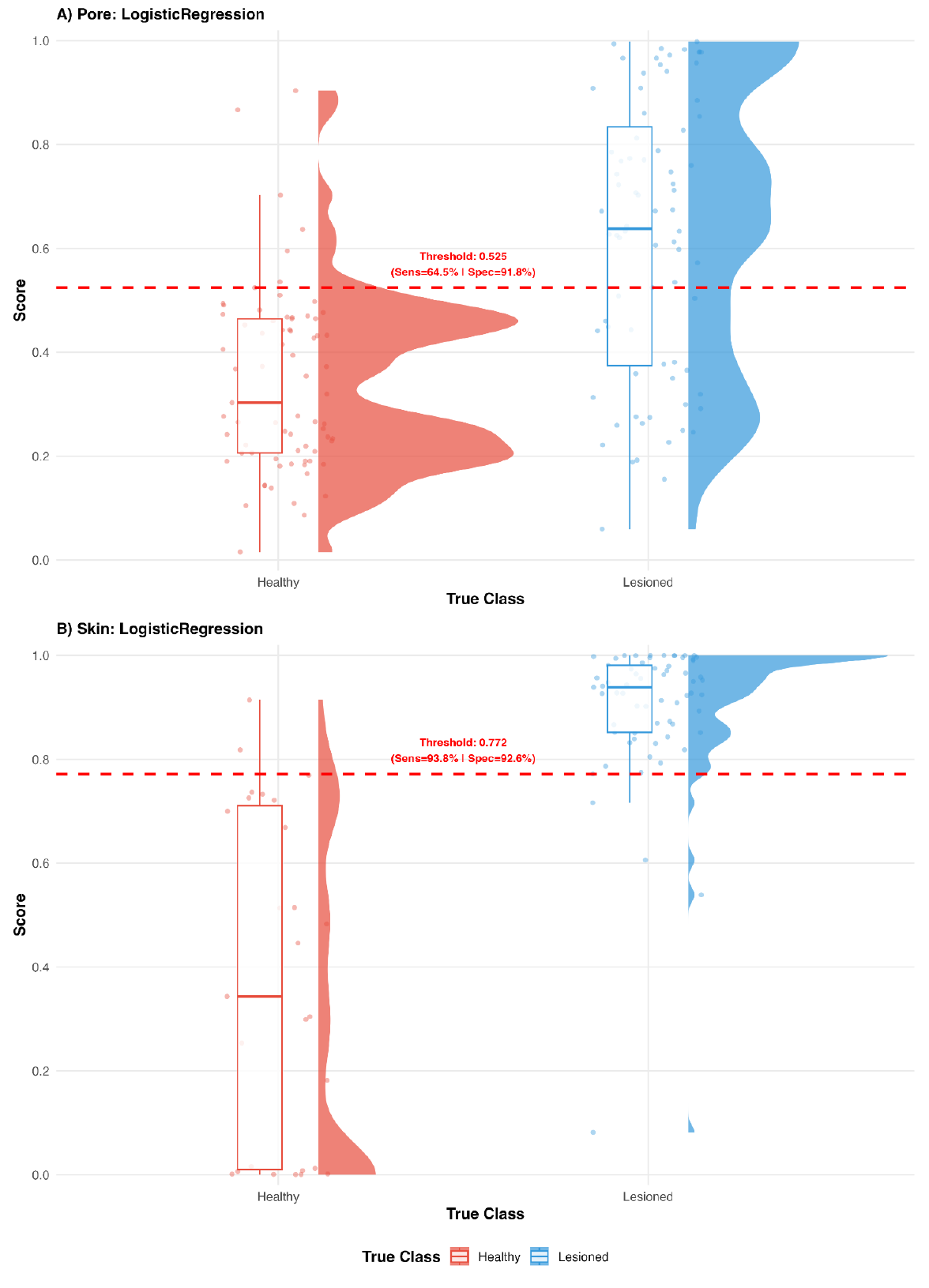
Microbiome Health Scores. Distribution of scores for the Logistic Regression model for A) urogenital pore samples, and B) skin samples. Each point represents an individual fish, colored by its clinical status (Red = healthy; Blue = Lesioned). While the models show clear separation between groups, individuals in the “transition zone” (scores 0.5–0.6 for pore and 0.7-0.8 for skin) represent microbial profiles that have diverged from the healthy baseline but may not yet exhibit severe clinical symptoms. This score provides a quantitative, continuous metric for health monitoring beyond binary visual diagnosis.

## Discussion

The transition toward sustainable aquaculture necessitates a paradigm shift: moving from reactive disease management to proactive monitoring based on the health of the microbial ecosystem. This study demonstrates that the fish microbiome is not merely a biological bystander, but a sensitive diagnostic asset capable of reflecting the host’s physiological state through quantifiable microbial signatures

Analysis of alpha diversity revealed no statistically significant differences in community structure across the tissue-status interaction, nor within the main effects of tissue or status. The Shannon and Simpson indices remained stable when comparing healthy and lesioned individuals, contrasting with the diversity loss narrative often associated with dysbiosis. This suggests a process of microbial replacement rather than simple community collapse, where specific taxa shift while overall richness and evenness are maintained. This finding aligns with the concept that stress leads to increased stochastic variation and community heterogeneity rather than a single directional shift (Ma, 2020; Zaneveld et al., 2017). The significantly higher dispersion observed in the gills of lesioned fish (p = 0.029) confirms this breakdown of homeostasis, providing direct support for the Anna Karenina principle in a salmonid aquaculture context Consequently, health may be better defined by the stability of conserved functional pathways rather than fixed taxonomic checklists (Mougin and Joyce, 2022).

A central finding of this study is the strong statistical coupling between external and internal mucosal sites. The Procrustes correlation coefficient (*r*_*Proc*_) and Mantel correlation (*r*_*M*_) between skin and gills (*r*_*Proc*_ = 0.84, *r*_*M*_ = 0.52; P < 0.001) positions skin swabs as a robust, non-invasive proxy for branchial health. Similarly, the urogenital-gut coupling (*r*_*Proc*_ = 0.70, *r*_*M*_ = 0.54; P < 0.001) confirms that external low-biomass sites accurately reflect the internal digestive state, with Procrustes analysis demonstrating a higher sensitivity to the shared community structure across tissues. It should be noted, however, that the skin and urogenital pore microbiomes may also independently encode health information, not solely through their relationship with internal sites. The machine learning models were trained directly on data from these external sites, and their classification performance validates the diagnostic utility of non-invasive sampling regardless of the precise mechanism of compositional coupling. Notably, the skin model achieved substantially higher performance (AUC = 0.967) than the urogenital pore model (AUC = 0.869), suggesting that the skin microbiome encodes a stronger signal of systemic health status in this cohort. This difference may reflect the more direct exposure of the skin surface to the external pathogens associated with the clinical signs used for classification (skin lesions, gill pallor).

The validation of these minimally invasive techniques is critical for animal welfare. Recent research confirms that external mucus can reflect internal conditions like chronic enteritis (Legrand et al., 2017) without compromising the fish’s barrier function (Sanahuja et al., 2023). This approach enables high-frequency monitoring and reduces the need for lethal sampling, aligning with modern sustainable diagnostic protocols (Zarantonello et al., 2024; (Clinton et al., 2021) further demonstrated that gill swabs isolate diverse microbial consortia with minimal host DNA contamination, supporting the practical feasibility of swab-based monitoring.

While traditional metrics like alpha diversity suggest stability, our machine learning results demonstrate that disease signals exist at deeper levels of compositional complexity. The high accuracy in our benchmarked Logistic Regression models demonstrates that microbiome profiles contain a reliable signature of health status. This is consistent with recent work showing that gut microbiome models can predict compromised environments with >90% accuracy regardless of geolocation (Turner et al., 2022). Bayesian networks have also been used to model how farming conditions alter diversity, allowing for pathogen suppression (Soriano et al., 2023).

To bridge the gap between complex bioinformatics and farm-level decision-making, we distilled these high-dimensional datasets into continuous, quantitative Score Indices, which moves health assessment beyond binary classifications. This approach parallels the development of the Gut Microbiome Wellness Index (GMWI) (Chang et al., 2024) in human medicine, where composite microbiome-based scores have proven effective for disease-agnostic health prediction. To our knowledge, the Salmon Microbiome Health Scores presented here represent the first formalized, multi-parameter microbiome health index for any farmed fish species, addressing a gap explicitly noted in recent reviews (Mougin and Joyce, 2022; Xavier et al., 2024).

The value of these indices lies in their continuous nature, which in principle could allow longitudinal tracking of physiological resilience. A rising score could serve as an early warning signal and provides a standardized endpoint to evaluate interventions such as functional feeds. This approach parallels human research, where biomarkers indicate disease development prior to physical evidence (Dong-Min et al., 2024).

### Limitations and future directions

Some limitations should be considered when interpreting these results. The study employs a cross-sectional design in which the ground truth for ML model training was clinical visual assessment. Consequently, the health scores classify the same health states that clinical examination identifies, albeit through a standardizable, quantitative metric. The scores cannot, by construction, be demonstrated to detect disease before clinical signs appear; validating true pre-symptomatic predictive capability would require longitudinal studies with serial sampling before disease onset. However, the identification of a “transition zone” (scores 0.4-0.6) in which fish show microbial profiles diverging from the healthy baseline suggests that intermediate states exist, and future temporal studies could determine whether these states precede overt disease.

All fish originated from a single production cohort at one commercial facility. Water microbiome, feed composition, host genetics, temperature, and husbandry practices all shape baseline microbial communities (“Environmental and physiological factors shape the gut microbiota of Atlantic salmon parr (Salmo salar L.),” 2017; Uren Webster et al., 2018), and the health score thresholds established here may not transfer directly to other farms or production systems without recalibration. Multi-farm validation across diverse geographic regions and production stages (freshwater, smolt, marine) is a critical next step.

The lesioned cohort included fish presenting heterogeneous clinical signs (skin ulcers, exophthalmos, pale gills), potentially associated with different underlying pathologies. While the disease-agnostic approach is intentional and mirrors the GMWI framework in human medicine, the relative contribution of specific diseases to the microbiome signal remains to be characterized. The absence of confirmatory diagnosis at this step (e.g. qPCR for specific pathogens) means that the specific etiological agents cannot be confirmed.

The loss of healthy skin samples during quality filtering, introduces a potential bias in the skin health score. If healthy fish routinely yield lower skin microbial mass, the practical applicability of skin-based monitoring may be constrained by sampling limitations in the target populations.

Finally, the outer cross-validation loop employed only two folds (group-blocked by sea cage), yielding only two independent performance estimates. While this design prevents data leakage between cage environments, the resulting performance metrics should be interpreted with appropriate caution regarding their precision.

Despite these limitations, the identified proxy relationship and classification performance establish a strong proof-of-concept. Future research should prioritize: (1) longitudinal studies to determine whether microbiome score changes precede clinical manifestation; (2) multi-farm validation to establish transferable thresholds; (3) integration with metagenomic or metatranscriptomics approaches to reveal functional states beyond taxonomic composition (Mougin and Joyce, 2022; Vera-Ponce de León et al., 2024); and (4) coupling with host genomic data to explore microbiome-host interactions influencing disease resistance (Nguyen, 2024). This diagnostic framework also provides a foundation for targeted microbiome engineering as an alternative to antibiotics. Probiotic interventions and precision-engineered synthetic microbial communities offer promising avenues for modulating host phenotypes (Tayyab et al., 2025), potentially guided by the taxa identified as health score drivers.

## Conclusions

This study demonstrates that the microbiome profiles of Atlantic salmon encode health status information that can be captured through non-invasive sampling and translated into quantitative health scores using machine learning. The statistical validation of the urogenital pore and skin as compositional proxies for internal mucosal environments enables high-frequency monitoring while minimizing animal stress. The two Salmon Microbiome Health Scores developed here, represent, to our knowledge, the first formalized multi-parameter microbiome health indices for a farmed fish species. While longitudinal validation is necessary to determine whether microbiome score changes precede clinical disease, the current framework establishes a foundation for a shift from reactive to proactive, microbiome-informed aquaculture health management, with the potential to reduce antibiotic dependence and improve aquaculture welfare.

## Acknowledgments

The authors would like to thank the veterinarians and staff at the commercial marine aquaculture facility for their assistance during sample collection.

## Author Contributions

LEL conceptualized the study. LEL, FF and AP developed the methodology. LEL, FF, and AP were responsible for software development and formal analysis of the microbiome data. CL, MIO, and AP conducted the investigation and sample collection at the marine facility. AP, DG, and AB provided resources and performed data curation. LEL and JAU wrote the original draft. All authors (LEL, CL, FF, AP, MIO, DG, JAU, and AB) contributed to writing, review, and editing of the final manuscript. LEL, and FF created the visualizations. AB and DG were responsible for funding acquisition

## References

Anderson, M.J., 2006. Distance-based tests for homogeneity of multivariate dispersions. Biometrics 62. 10.1111/j.1541-0420.2005.00440.x

Auclert, L.Z., Chhanda, M.S., Derome, N., 2024. Interwoven processes in fish development: microbial community succession and immune maturation. PeerJ 12. 10.7717/peerj.17051

Bozzi, D., Rasmussen, J.A., Carøe, C., Sveier, H., Nordøy, K., Gilbert, M.T.P., Limborg, M.T., 2021. Salmon gut microbiota correlates with disease infection status: potential for monitoring health in farmed animals. Animal microbiome 3. 10.1186/s42523-021-00096-2

Callahan, B.J., McMurdie, P.J., Rosen, M.J., Han, A.W., Johnson, A.J., Holmes, S.P., 2016. DADA2: High-resolution sample inference from Illumina amplicon data. Nature methods 13. 10.1038/nmeth.3869

Chang, D., Gupta, V.K., Hur, B., Cobo-López, S., Cunningham, K.Y., Han, N.S., Lee, I., Kronzer, V.L., Teigen, L.M., Karnatovskaia, L.V., Longbrake, E.E., Davis, J.M., Nelson, H., Sung, J., 2024. Gut Microbiome Wellness Index 2 enhances health status prediction from gut microbiome taxonomic profiles. Nature communications 15. 10.1038/s41467-024-51651-9

Clinton, M., Wyness, A.J., Martin, S.A.M., Brierley, A.S., Ferrier, D.E.K., 2024. Association of microbial community structure with gill disease in marine-stage farmed Atlantic salmon (Salmo salar); a yearlong study. BMC Veterinary Research 20, 340.

Clinton, M., Wyness, A.J., Martin, S.A.M., Brierley, A.S., Ferrier, D.E.K., 2021. Sampling the fish gill microbiome: a comparison of tissue biopsies and swabs. BMC microbiology 21. 10.1186/s12866-021-02374-0

Dehler, C.E., Secombes, C.J., Martin, S.A.M., 2017. Seawater transfer alters the intestinal microbiota profiles of Atlantic salmon (Salmo salar L.). Scientific reports 7. 10.1038/s41598-017-13249-8

Dong-Min, J., Morton, J.T., Bonneau, R., 2024. Meta-analysis of the human gut microbiome uncovers shared and distinct microbial signatures between diseases. mSystems 9. 10.1128/msystems.00295-24

Environmental and physiological factors shape the gut microbiota of Atlantic salmon parr (Salmo salar L.), 2017 Aquaculture 467, 149–157.

Evensen, Ø., 2016. Immunization Strategies against Piscirickettsia salmonis Infections: Review of Vaccination Approaches and Modalities and Their Associated Immune Response Profiles. Frontiers in immunology 7. 10.3389/fimmu.2016.00482

Froehlich, H.E., Gentry, R.R., Halpern, B.S., 2018. Global change in marine aquaculture production potential under climate change. Nature ecology & evolution 2. 10.1038/s41559-018-0669-1

Gajardo, K., Rodiles, A., Kortner, T.M., Krogdahl, Å., Bakke, A.M., Merrifield, D.L., Sørum, H., 2016. A high-resolution map of the gut microbiota in Atlantic salmon (Salmo salar): A basis for comparative gut microbial research. Scientific reports 6. 10.1038/srep30893

Hajjo, R., Sabbah, D.A., Al Bawab, A.Q., 2022. Unlocking the Potential of the Human Microbiome for Identifying Disease Diagnostic Biomarkers. Diagnostics (Basel, Switzerland) 12. 10.3390/diagnostics12071742

Klindworth, A., Pruesse, E., Schweer, T., Peplies, J., Quast, C., Horn, M., Glöckner, F.O., 2013. Evaluation of general 16S ribosomal RNA gene PCR primers for classical and next-generation sequencing-based diversity studies. Nucleic Acids Res 41, e1.

Legrand, T.P.R.A., Catalano, S.R., Wos-Oxley, M.L., Stephens, F., Landos, M., Bansemer, M.S., Stone, D.A.J., Qin, J.G., Oxley, A.P.A., 2017. The Inner Workings of the Outer Surface: Skin and Gill Microbiota as Indicators of Changing Gut Health in Yellowtail Kingfish. Front Microbiol 8, 2664.

Limborg, M.T., Alberdi, A., Kodama, M., Roggenbuck, M., Kristiansen, K., Gilbert, M.T.P., 2018. Applied Hologenomics: Feasibility and Potential in Aquaculture. Trends in biotechnology 36. 10.1016/j.tibtech.2017.12.006

Mabrok, M., Algammal, A.M., Sivaramasamy, E., Hetta, H.F., Atwah, B., Alghamdi, S., Fawzy, A., Avendaño-Herrera, R., Rodkhum, C., 2023. Tenacibaculosis caused by Tenacibaculum maritimum: Updated knowledge of this marine bacterial fish pathogen. Frontiers in cellular and infection microbiology 12. 10.3389/fcimb.2022.1068000

Ma, Z.S., 2020. Testing the Anna Karenina Principle in Human Microbiome-Associated Diseases. iScience 23. 10.1016/j.isci.2020.101007

McMurdie, P.J., Holmes, S., 2013. phyloseq: an R package for reproducible interactive analysis and graphics of microbiome census data. PloS one 8. 10.1371/journal.pone.0061217

Mitchell, S.O., Rodger, H.D., 2011. A review of infectious gill disease in marine salmonid fish. Journal of fish diseases 34. 10.1111/j.1365-2761.2011.01251.x

Mougin, J., Joyce, A., 2022. Fish disease prevention via microbial dysbiosis-associated biomarkers in aquaculture. Rev. Aquac. 10.1111/raq.12745

Nguyen, N.H., 2024. Genetics and Genomics of Infectious Diseases in Key Aquaculture Species. Biology 13. 10.3390/biology13010029

Noble, C., Abbink, W., Alvestad, R., Ardó, L., Bégout, M.-L., Bloecher, N., Burgerhout, E., Calduch-Giner, J., Chivite-Alcalde, M., Císař, P., Durland, E., Espmark, Å.M., Falconer, L., Føre, M., Georgopoulou, D.G., Heia, K., Helberg, G.A.N., Gomez, D.I., Johansen, L.-H., Johansson, G.S., Jónsdóttir, K.E., Kolarevic, J., Krasnov, A., Kumaran, S.K., Kvæstad, B., Larsson, T., Lazado, C.C., Madaro, A., Moroni, F., Måge, I., Nilsson, J., Ortega, S., Papandroulakis, N., Pérez-Sánchez, J., Prentice, P.M., Planellas, S.R., Roth, B., Smith, A., Solberg, L.E., Stavrakidis-Zachou, O., Stien, L.H., Striberny, A., Svalheim, R.A., Sæther, B.-S., Timmerhaus, G., Toften, H., Tschirren, L., van de Vis, H., Ytteborg, E., Zena, L.A., Østbye, T.-K.K., 2026. Welfare Indicators for Aquaculture Research: Toolboxes for Five Farmed European Fish Species. Reviews in Aquaculture 18, e70109.

Noble, C., Jones, H.A., Damsgård, B., Flood, M.J., Midling, K.Ø., Roque, A., Sæther, B.S., Cottee, S.Y., 2012. Injuries and deformities in fish: their potential impacts upon aquacultural production and welfare. Fish physiology and biochemistry 38. 10.1007/s10695-011-9557-1

Parks, D.H., Chuvochina, M., Rinke, C., Mussig, A.J., Chaumeil, P.A., Hugenholtz, P., 2022. GTDB: an ongoing census of bacterial and archaeal diversity through a phylogenetically consistent, rank normalized and complete genome-based taxonomy. Nucleic acids research 50. 10.1093/nar/gkab776

Peres-Neto, P.R., Jackson, D.A., 2001. How well do multivariate data sets match? The advantages of a Procrustean superimposition approach over the Mantel test. Oecologia 129. 10.1007/s004420100720

Roberts, D.R., Bahn, V., Ciuti, S., Boyce, M.S., Elith, J., Guillera-Arroita, G., Hauenstein, S., Lahoz-Monfort, J.J., Schröder, B., Thuiller, W., Warton, D.I., Wintle, B.A., Hartig, F., Dormann, C.F., 2017. Cross-validation strategies for data with temporal, spatial, hierarchical, or phylogenetic structure. Ecography 40, 913–929.

Rozas-Serri, M., 2022. Why Does Piscirickettsia salmonis Break the Immunological Paradigm in Farmed Salmon? Biological Context to Understand the Relative Control of Piscirickettsiosis. Frontiers in immunology 13. 10.3389/fimmu.2022.856896

Rozas-Serri, M., Kani, T., Jaramillo, V., Correa, R., Ildefonso, R., Rabascall, C., Barrientos, S., Coñuecar, D., Peña, A., 2024. Current vaccination strategy against Piscirickettsia salmonis in Chile based only on the EM-90 genogroup shows incomplete cross-protection for the LF-89 genogroup. Fish & shellfish immunology 154. 10.1016/j.fsi.2024.109893

Sanahuja, I., Guerreiro, P.M., Girons, A., Fernandez-Alacid, L., Ibarz, A., 2023. Evaluating the repetitive mucus extraction effects on mucus biomarkers, mucous cells, and the skin-barrier status in a marine fish model. Front. Mar. Sci. 9. 10.3389/fmars.2022.1095246

Soriano, B., Hafez, A.I., Naya-Català, F., Moroni, F., Moldovan, R.A., Toxqui-Rodríguez, S., Piazzon, M.C., Arnau, V., Llorens, C., Pérez-Sánchez, J., 2023. SAMBA: Structure-Learning of Aquaculture Microbiomes Using a Bayesian Approach. Genes (Basel) 14. 10.3390/genes14081650

Spilsberg, B., Nilsen, H.K., Tavornpanich, S., Gulla, S., Jansen Lagesen, K., Colquhoun, D.J., Olsen, A.B., 2022. Tenacibaculosis in Norwegian Atlantic salmon (Salmo salar) cage-farmed in cold sea water is primarily associated with Tenacibaculum finnmarkense genomovar finnmarkense. Journal of fish diseases 45. 10.1111/jfd.13577

Tayyab, M., Zhao, Y., Zhang, Y., 2025. Microbiome engineering to enhance disease resistance in aquaculture: current strategies and future directions. Front. Microbiol. 16, 1625265.

Topçuoğlu, B.D., Lesniak, N.A., Ruffin, M.T., Wiens, J., Schloss, P.D., 2020. A Framework for Effective Application of Machine Learning to Microbiome-Based Classification Problems. mBio 11. 10.1128/mBio.00434-20

Troell, M., Naylor, R.L., Metian, M., Beveridge, M., Tyedmers, P.H., Folke, C., Arrow, K.J., Barrett, S., Crépin, A.S., Ehrlich, P.R., Gren, A., Kautsky, N., Levin, S.A., Nyborg, K., Österblom, H., Polasky, S., Scheffer, M., Walker, B.H., Xepapadeas, T., de Zeeuw, A., 2014. Does aquaculture add resilience to the global food system? Proceedings of the National Academy of Sciences of the United States of America 111. 10.1073/pnas.1404067111

Turner, J.W., Cheng, X., Saferin, N., Yeo, J.Y., Yang, T., Joe, B., 2022. Gut microbiota of wild fish as reporters of compromised aquatic environments sleuthed through machine learning. Physiological genomics 54. 10.1152/physiolgenomics.00002.2022

Uren Webster, T.M., Consuegra, S., Hitchings, M., de Leaniz, C.G., 2018. Interpopulation Variation in the Atlantic Salmon Microbiome Reflects Environmental and Genetic Diversity. Applied and Environmental Microbiology. 10.1128/AEM.00691-18

Vera-Ponce de León, A., Hensen, T., Hoetzinger, M., Gupta, S., Weston, B., Johnsen, S.M., Rasmussen, J.A., Clausen, C.G., Pless, L., Veríssimo, A.R.A., Rudi, K., Snipen, L., Karlsen, C.R., Limborg, M.T., Bertilsson, S., Thiele, I., Hvidsten, T.R., Sandve, S.R., Pope, P.B., La Rosa, S.L., 2024. Genomic and functional characterization of the Atlantic salmon gut microbiome in relation to nutrition and health. Nature Microbiology 9, 3059–3074.

Wang, A.R., Ran, C., Ringø, E., Zhou, Z.G., 2018. Progress in fish gastrointestinal microbiota research. Reviews in Aquaculture 10, 626–640.

Wang, J., Li, Y., Jaramillo-Torres, A., Einen, O., Jakobsen, J.V., Krogdahl, Å., Kortner, T.M., 2023. Exploring gut microbiota in adult Atlantic salmon (Salmo salar L.): Associations with gut health and dietary prebiotics. Animal microbiome 5. 10.1186/s42523-023-00269-1

Wynne, J.W., Thakur, K.K., Slinger, J., Samsing, F., Milligan, B., Powell, J.F.F., McKinnon, A., Nekouei, O., New, D., Richmond, Z., Gardner, I., Siah, A., 2020. Microbiome Profiling Reveals a Microbial Dysbiosis During a Natural Outbreak of Tenacibaculosis (Yellow Mouth) in Atlantic Salmon. Front. Microbiol. 11, 586387.

Xavier, R., Severino, R., Silva, S.M., 2024. Signatures of dysbiosis in fish microbiomes in the context of aquaculture. Reviews in Aquaculture 16, 706–731.

Xiong, J.B., Nie, L., Chen, J., 2019. Current understanding on the roles of gut microbiota in fish disease and immunity. Zoological research 40. 10.24272/j.issn.2095-8137.2018.069

Zaneveld, J.R., McMinds, R., Vega, T.R., 2017. Stress and stability: applying the Anna Karenina principle to animal microbiomes. Nature microbiology 2. 10.1038/nmicrobiol.2017.121

